# Prediction and Evolution of B Cell Epitopes of Surface Protein in SARS-CoV-2

**DOI:** 10.1101/2020.04.03.022723

**Authors:** Jerome R Lon, Yunmeng Bai, Bingxu Zhong, Fuqaing Cai, Hongli Du

## Abstract

The discovery of epitopes is helpful to the development of SARS-CoV-2 vaccine. The sequences of the surface protein of SARS-CoV-2 and its proximal sequences were obtained by BLAST, the sequences of the whole genome of SARS-CoV-2 were obtained from the GenBank. Based on the NCBI Reference Sequence: NC_045512.2, the conformational and linear B cell epitopes of the surface protein were predicted separately by various prediction methods. Furthermore, the conservation of the epitopes, the adaptability and other evolutionary characteristics were also analyzed. 7 epitopes were predicted, including 5 linear epitopes and 2 conformational epitopes, one of the linear and one of the conformational were coincide. The epitope D mutated easily, but the other epitopes were very conservative and the epitope C was the most conservative. It is worth mentioning that all of the 6 dominated epitopes were absolutely conservative in nearly 1000 SARS-CoV-2 genomes, and they deserved further study. The findings would facilitate the vaccine development, had the potential to be directly applied on the treatment in this disease, but also have the potential to prevent the possible threats caused by other types of coronavirus.

## Introduction

In late December 2019, a novel coronavirus was officially named as SARS-CoV-2 by World Health Organization(WHO) and identified as the pathogen causing outbreaks of SARS-like and MERS-like illness in Chinese city of Wuhan, which was a zoonotic disease. As of March 13, 2020, the outbreak of SARS-CoV-2 has been reported in many areas of the world, with more than 130,000 people infected[1]. With an alarmingly human-to-human transmissibility, the reproductive number of SARS-CoV-2 has been computed to around 3.28 [2]. According to the data in NGDC(National Genomics Data Center), at 15:00(GMT+8) on March 13, 2020, 482 genomic variations of SARS-CoV-2 has been reported, which has aroused widespread concern.

The B cell epitope of viral surface protein can specifically bind to the host’s B cell antigen receptor and induce the body to produce protective antibody and humoral immune response. The discovery of epitopes is helpful to the development of SARS-CoV-2 vaccine and the understanding of SARS-CoV-2’s pathogenesis[3]. 3 proteins embedded in the virus envelope of SARS-CoV-2 have been identified, Spike(S) protein, Envelope(E) protein, Membrane(M) protein. At present, due to the lack of study of the crystal structure of surface protein of SARS-CoV-2, the study of epitopes is time-consuming, power-consuming, cost and difficult [4].

In this work, we analyzed the surface protein of SARS-CoV-2, predicted the structures with bioinformatics methods. On the basis, we predicted the linear and conformational B cell epitopes, analyzed the conservation of the epitopes, the adaptability and other evolutionary characteristics of the surface protein, which provided a theoretical basis for the vaccine development and prevention of SARS-CoV-2.

## Results

### Basic analysis of surface protein of SARS-CoV-2

The primary structure and physicochemical properties of the S/E/M protein were analyzed. The results revealed that the S protein has an average hydrophilic index of −0.079**(Figure S1A)**. On the basis of hydrophilicity, it also showed amphoteric properties. There was an outside-in transmembrane helix in 23 residues from position 1214^th^ to position 1236^th^ at the N-terminal(Figure S2A). The protein instability index was 33.01, which revealed the S protein was stable. The E protein has an average hydrophilic index of 1.128(Figure S1B). It was hydrophobic. An inside-out transmembrane helix in 23 residues from position 12^th^ to position 34^th^ at the N-terminal was predicted(Figure S2B). The protein instability index was 38.68, which revealed the E protein was stable. The M protein has an average hydrophilic index of 0.446(Figure S1C). On the basis of hydrophobicity, it also showed amphoteric properties. There were two outside-in transmembrane helices, one was in 20 residues from position 20^th^ to position 39^th^, the another one was in 23 residues from position 78^th^ to position 100^th^, and an inside-out transmembrane helix in 20 residues from position 51^st^ to position 73^rd^, at the N-terminal(Figure S2C). The protein instability index was 39.14, which revealed the M protein was stable.

### Prediction of the 3D structure of surface protein of SARS-CoV-2

The optimal template for homology modeling of the S protein of SARS-CoV-2 was the S protein of SARS(PDB ID: 6acc.1), with the sequence identity of 76.47% and the GMQE score of 0.73. According to the evaluation of the structure by Ramachandran plot(**Figure 1A**), 99.3% of the residues were located in the most favoured regions and the allowed regions, 0.7% of the residues were located in the disallowed regions, the high-energy regions(**Table 1**), which was possibly due to some energy was spent in the protein processing to make these residues enter the high-energy regions [5]. The result generally showed that the structure was reliable. The structure of S protein(Figure 1B) of SARS-CoV-2 is a trimer, which can be divided into a tightly curled tail and a distributed head. The head is mainly composed of β-sheet, irregular curl and turn, which is exposed to the envelope of the virus, contributes to the formation of epitopes. The tail is mainly composed of several α-helices, part of which is embedded in the envelope, hinders the formation of epitopes.

**Table 1.** The plot statistics of the Ramachandran plot.

| Plot statistics-S |  |  | Plot statistics-E |  |  |
| --- | --- | --- | --- | --- | --- |
| Residues in most favoured regions [A, B, L] | 2352 | 78.80% | Residues in most favoured regions [A, B, L] | 228 | 84.40% |
| Residues in additional allowed regions [a, b, l, p] | 577 | 19.30% | Residues in additional allowed regions [a, b, l, p] | 38 | 14.10% |
| Residues in generously allowed regions [∼a, ∼b, ∼l, ∼p] | 36 | 1.20% | Residues in generously allowed regions [∼a, ∼b, ∼l, ∼p] | 4 | 1.50% |
| Residues in disallowed regions | 20 | 0.70% | Residues in disallowed regions | 0 | 0.00% |

**Figure 1.**
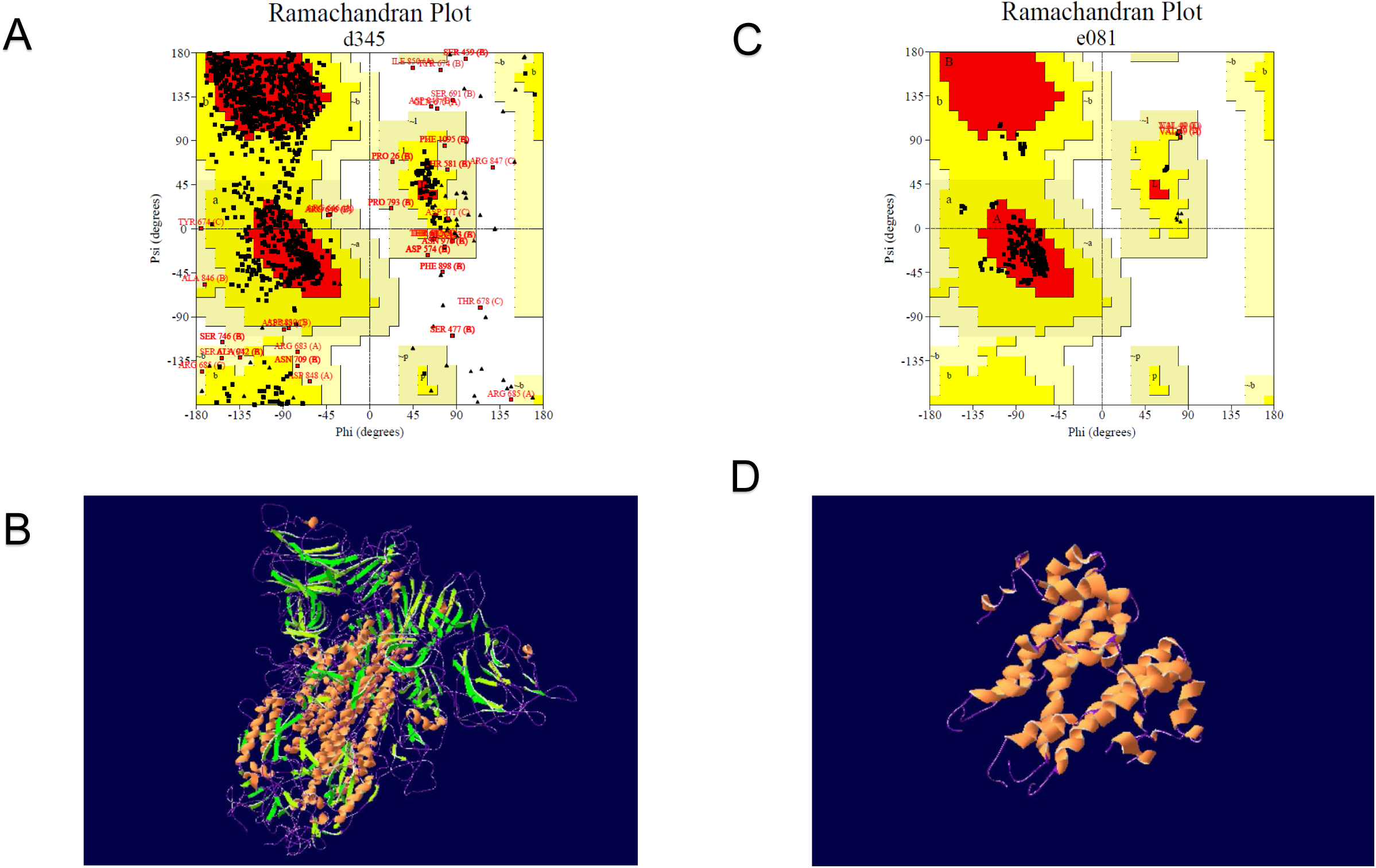
The 3D structure prediction and Ramachandran plot analysis of the S and E protein. **A.** The 3D structure of the S protein predicted by homology modeling. It is a trimer, the head contains RBD (receptor Binding Domain) [24], the tail contains the basic elements required for the membrane fusion, the end of the tail is a transmembrane region and is embedded in the envelope of SARS-CoV-2. **B.** The 3D structure of the E protein predicted by homology modeling. It is a pentamer with ion channel activity [25]. Its head is short, the middle of the tail is a transmembrane region which help the E protein embed in the envelope of SARS-CoV-2. **C.** The Ramachandran plot analysis of the 3D structure of the S protein (without Gly and Pro). Most residues located in the red (core) regions, and few in the white regions. **D.** The Ramachandran plot analysis of the 3D structure of the E protein (without Gly and Pro). All of the residues located on the red(core) region.

The optimal template for homology modeling of the E protein of SARS-CoV-2 was the E protein of SARS(PDB ID: 5×29.1), with the sequence identity of 91.38% and the GMQE score of 0.73. According to the evaluation of the structure by Ramachandran plot(Figure 1C), 100% of the residues were located in the most favoured regions(Table 1), indicating that the structure was reliable. The E protein of SARS-CoV-2 is a pentamer(Figure 1D), which can be divided into the concentrated transmembrane part and the head located outside the envelope. The head is mainly composed of α-helix, irregular curl and turn, which is exposed to the envelope, contributes to the formation of epitopes. The tail is mainly composed of long α-helix, most of which are embedded in the envelope, hinders the formation of epitopes.

The optimal template for homology modeling of the M protein of SARS-CoV-2 was the effector protein Zt-KP6-1(PDB ID: 6qpk. 1. A), with the sequence identity of 20.00% and the GMQE score of 0.06. The sequence identity between the optimal template and the M protein of SARS-CoV-2 and the GMQE score are too low, so that the template is not suitable for homology modeling.

### Prediction of linear B cell epitopes

All linear B cell epitopes of the surface protein were filtered according to the following criteria: (1) region with high surface probability(≥0.75), strong antigenicity(≥0) and high flexibility; (2) excluding the region with α-helix, β-sheet and glycosylation site(Figure 2); (3) in line with the prediction by BepiPred 2.0(cut off to 0.35)**(Table S1)** and ABCpred(cut off to 0.51)(Table S2). Based on the results obtained with these methods and artificial optimization, 4 potential linear B cell epitopes of the S protein were predicted(Table 2, Figure 3A), including 601-605 aa, 656-660 aa, 676-682 aa, 808-813 aa, and they were named as the epitope A, B, C, D, respectively; 1 epitope of the E protein was selected(60-65 aa) and named as the epitope F(Table 2, Figure 3C); 1 epitope of the M protein was selected (211-215 aa) and named as the epitope H(Table 2).

**Table 2.**
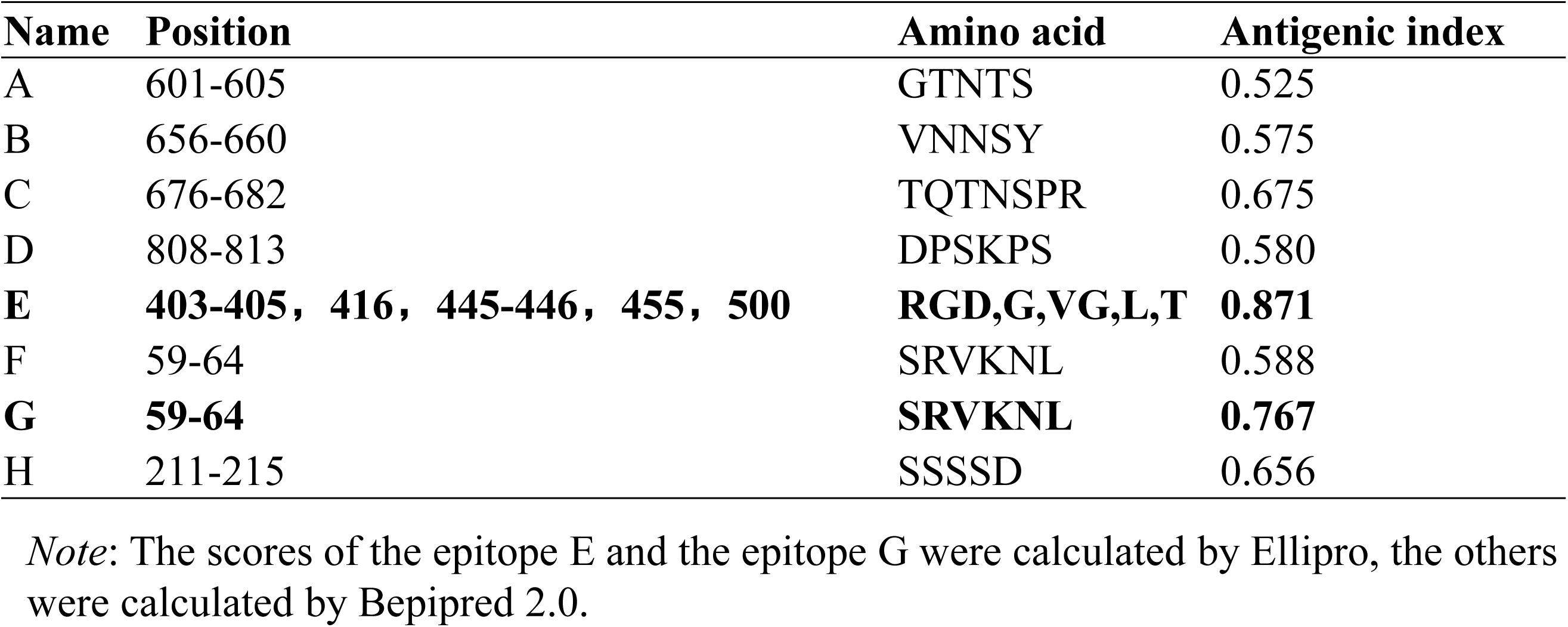
The composition and the antigenic index of the epitopes of SARS-CoV-2.

| **Name** | **Position** | **Amino acid** | **Antigenic index** |
| --- | --- | --- | --- |
| A | 601-605 | GTNTS | 0.525 |
| B | 656-660 | VNNSY | 0.575 |
| C | 676-682 | TQTNSPR | 0.675 |
| D | 808-813 | DPSKPS | 0.580 |
| **E** | **403-405, 416, 445-446, 455, 500** | **RGD,G,VG,L,T** | **0.871** |
| F | 59-64 | SRVKNL | 0.588 |
| **G** | **59-64** | **SRVKNL** | **0.767** |
| H | 211-215 | SSSSD | 0.656 |

*Note:* The scores of the epitope E and the epitope G were calculated by Ellipro, the others were calculated by Bepipred 2.0.

**Figure 2.**
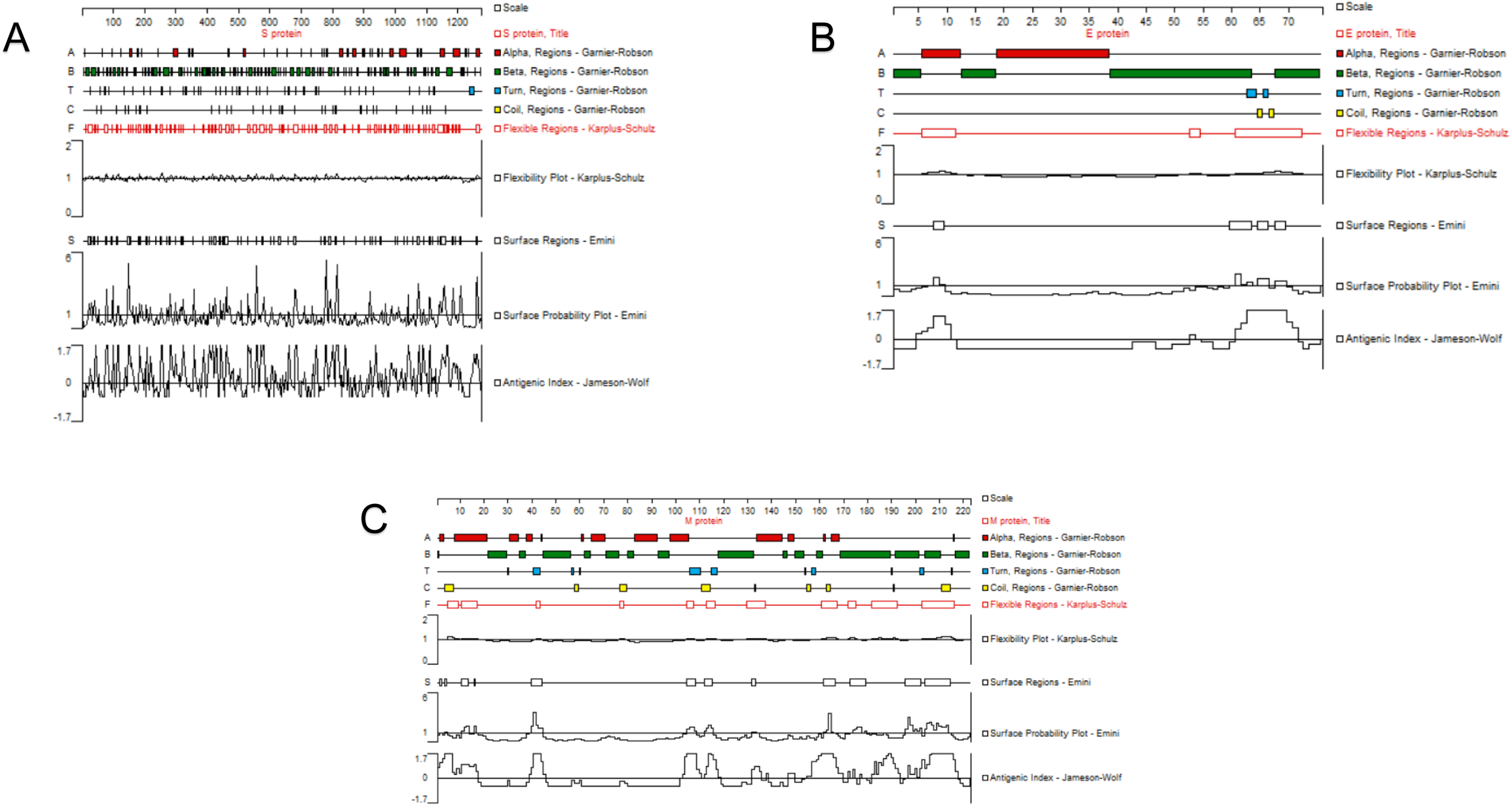
The secondary structures and properties analysis of the S, E and M protein with the Protean tool of DNAStar. **A.** Analysis of the S protein. It contains most α-helix and β-sheet, some Turn and Coli region, several discontinuous high flexibility fragments, fluctuant surface probablity with a few of positive peak and several antigenicity regions with positive peak. **B.** Analysis of the E protein. It contains most α-helix and β-sheet, some Turn and Coli region, three high flexibility fragments, few surface probablity regions and two antigenicity regions with positive peak in the begin and the end of polypeptide chain, respectively. **C.** Analysis of the M protein. It contains most α-helix and β-sheet, some Turn and Coli region, several high flexibility fragments, few surface probablity regions, two antigenicity regiona with positive single peak in the begin and middle of peptide chain, respectively, and consecutive positive peaks in the end.

**Figure 3.**
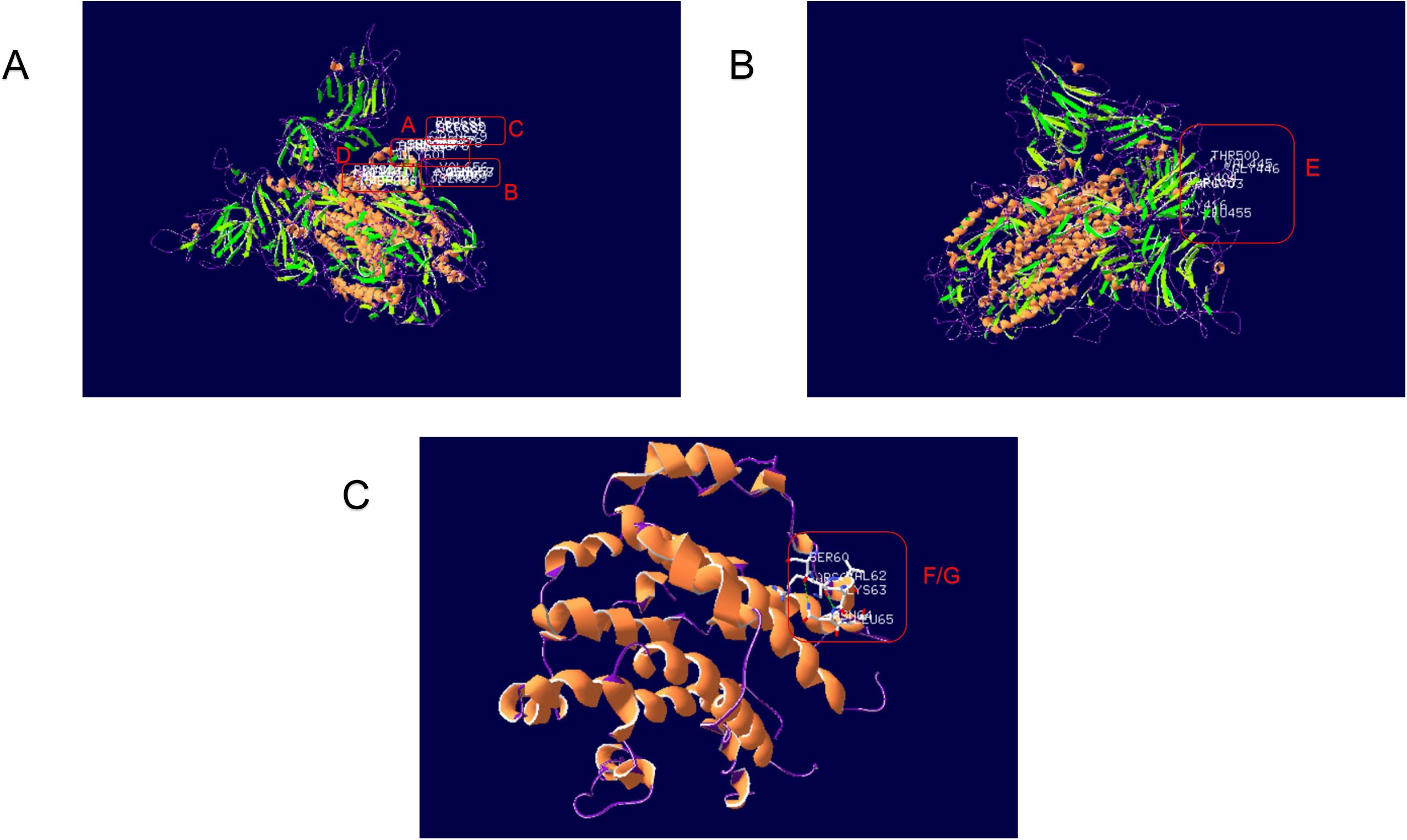
The predicted epitopes of the S and E protein. **A.** The predicted linear B-cell epitopes of the S protein. The epitope A, B, C located in the forepart of the tail, the epitope D located in the back part of the tail and is close to the transmembrane region. **B.** The predicted conformational B-cell epitope of the S protein. It located in the RBD of the head which is the vital sites binding with ACE2. **C.** The predicted B-cell epitope of the E protein. The epitope G is the linear epitope and the F is the conformational epitope, which are coincide.

### Prediction of conformational B cell epitopes

The conformational B cell epitopes of surface protein were predicted with Ellipro(Table S3) and SEPPA 3.0(Table S4) with the threshold of 0.063 and 0.5, respectively. After the artificial optimization, one conformational B-cell epitope (403-405,416,445,446,455,500 aa) of S protein was predicted(Table 2). It is obvious that the region located on the head of the S protein(Figure 3B), which is the outside of SARS-CoV-2, making it easy to form an epitope. We selected it as a dominant conformational epitope and named it as the epitope E. Additionally, one conformational B-cell epitope(60-65 aa) of E protein was predicted(Table 2), which is consistent with the epitope F of the E protein. Similarly, this region located on the outside(Figure 3C), we selected it as a dominant conformational epitope and named G. However, the conformational epitope of the M protein could not be predicted due to the failure of credible homology modeling.

### Analysis of epitope conservation

The Consurf Server was used to predict epitope conservative sites with the structure of surface proteins and the alignment results in different datasets (Table S5). All the epitopes of the S, E, M protein were absolute conservative among all SARS-CoV-2 sequences(Table 3A, Figure S3A-G). To further calculate the conservation of the epitopes in different coronavirus datasets, the representative sequences from SARS-CoV-2 were selected to participate in the human coronavirus dataset and the coronavirus dataset, due to amino acid sequences of some S or E or M protein were absolute conservative in SARS-CoV-2. The conservation was a little lower in human coronavirus than those of in SARS-CoV-2(Table 3B, Figure S4A-G), and the epitope D was easy to mutate. The other epitopes were conservative and the epitope F/G was the most conservative one. As for the coronavirus(Table 3C, Figure S5A-G), in the 5 epitopes of S protein, 4 of which obtained the conservative score less than 1, ranging from −0.854 to 0.256. It showed that the epitope C with the minimum score is the most stable and not easy to mutate. Besides, the score of the epitope D was 1.247, showing the relatively high possibility to mutate. The epitopes of the E protein were stable with the conservative score less than 1. The site conservation of the M protein could not be predicted due to the failure of credible homology modeling.

**Table 3A.**
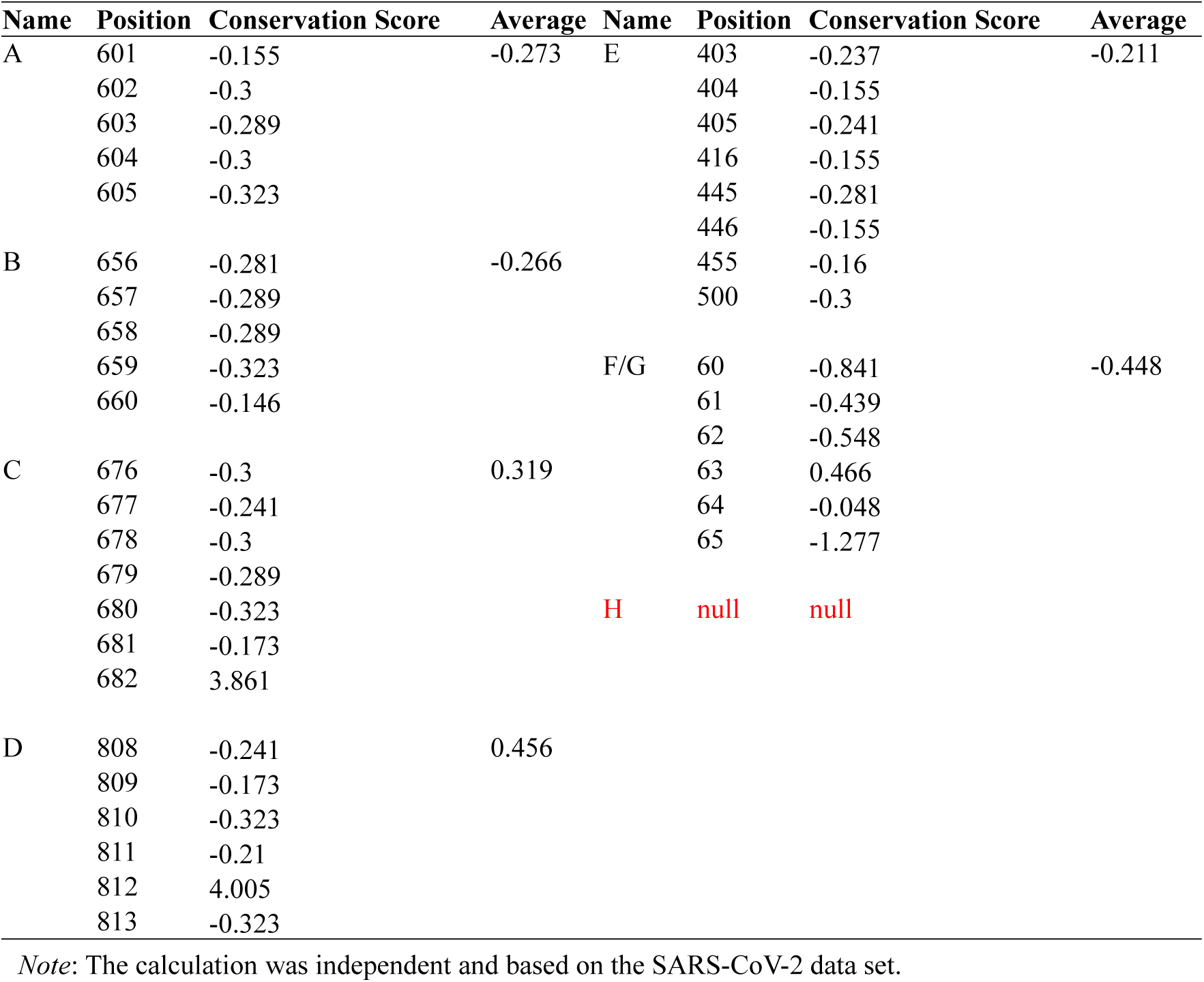
The conservation of the epitopes in SARS-CoV-2.

**Table 3B.**
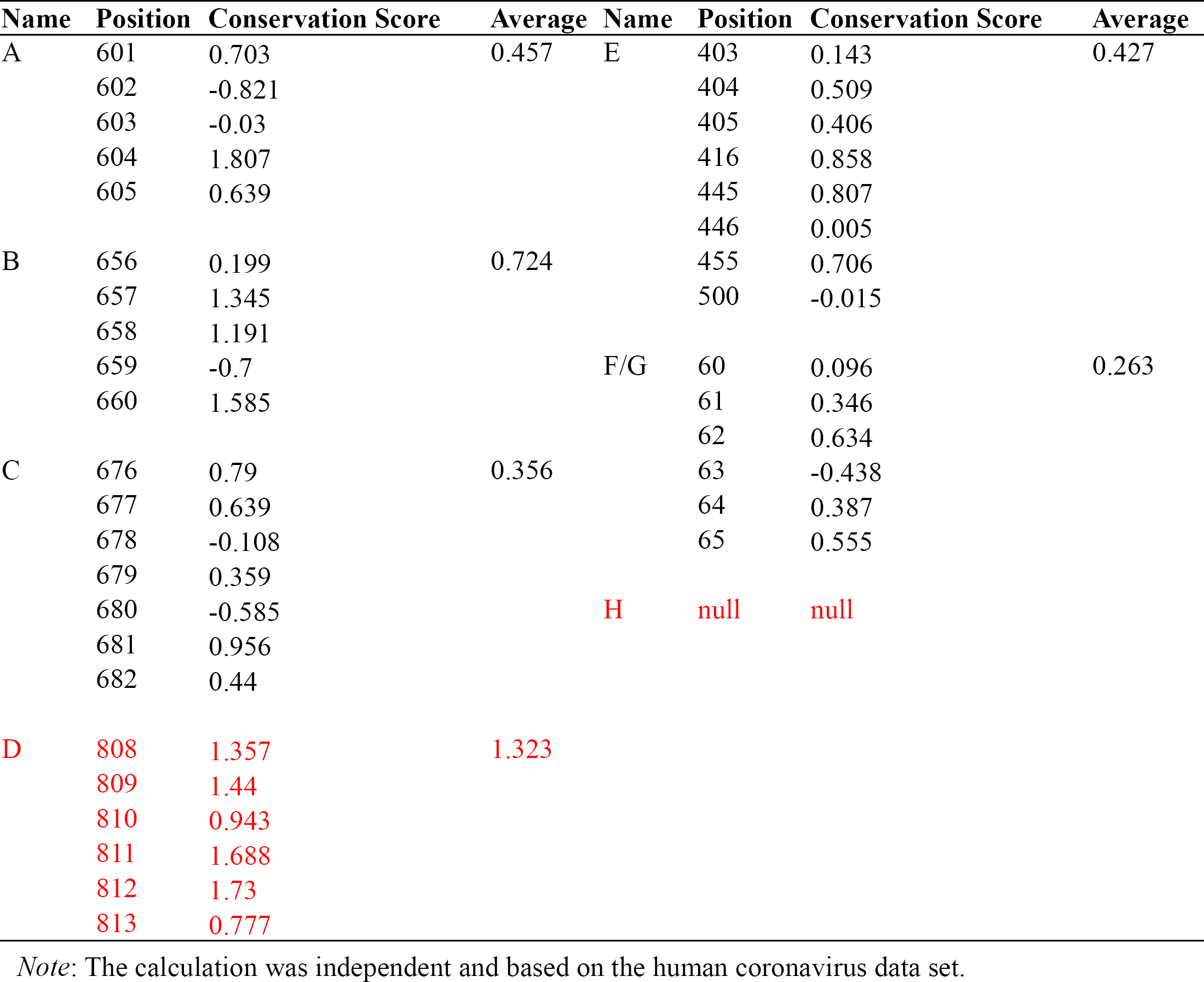
The conservation of the epitopes in human coronavirus.

**Table 3C.**
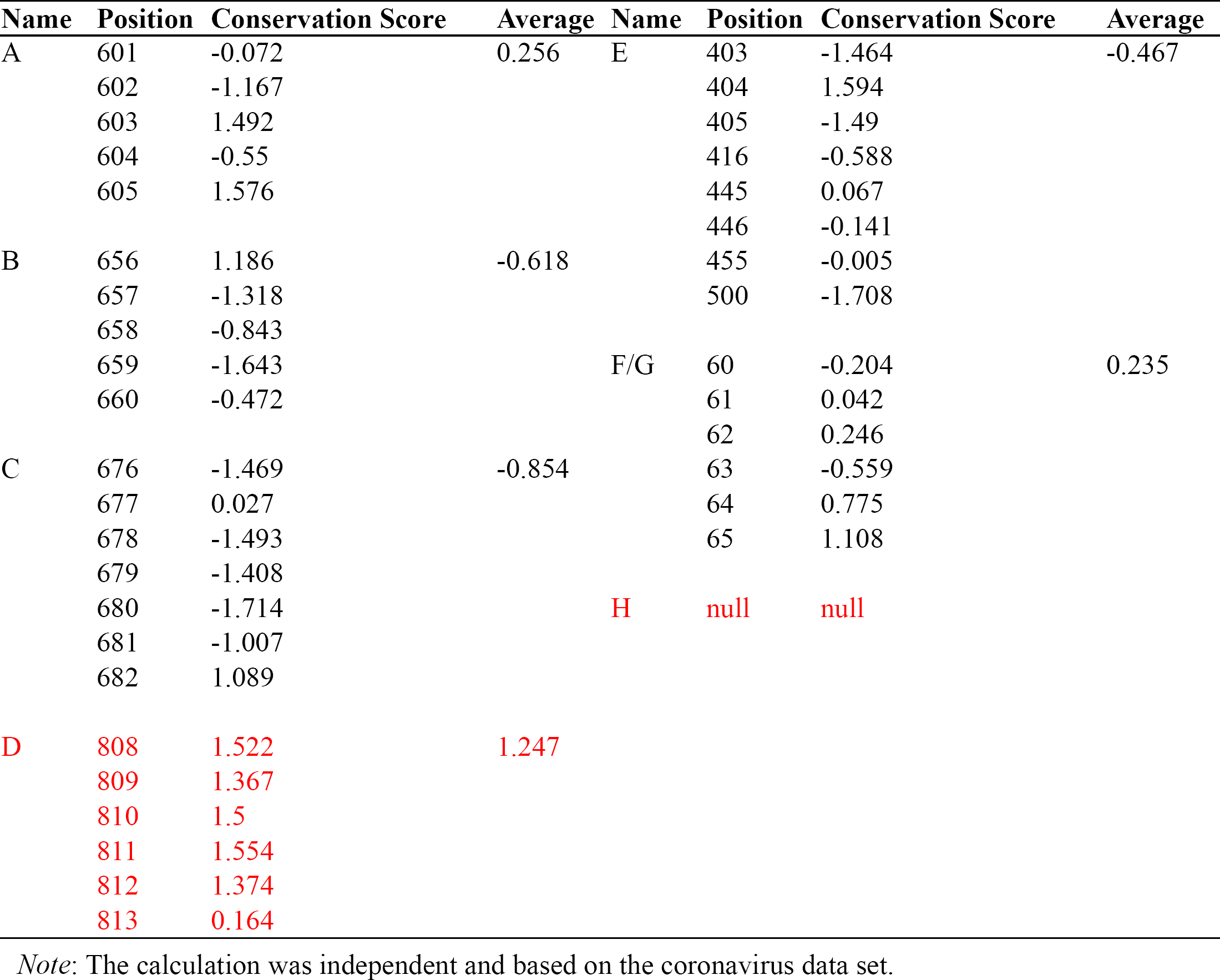
The conservation of the epitopes in coronavirus.

## Discussion

SARS-CoV-2 caused huge impact to human production, living and even life, has become a major challenge confronting the whole world. Development of vaccine is one of the effective means of prevention and treatment of the virus long-term. Epitope vaccine is the trend of development of vaccine due to the advantages of strong pertinence, less toxic and side effects and easy to transportation and storage [6]. The determination of epitopes is the basis of the development and application of vaccine, and the clinical diagnosis and treatment. Currently, the methods which were mainly used are X-crystal diffraction method, immune experiment method and bioinformatics method. The first two are time-consuming and laborious, the bioinformatics method is gaining more and more credibility among researchers [3,6,7]. There are many factors to be considered in the prediction of epitopes by bioinformatics method, such as the surface probablity and flexibility of the epitopes. At the same time, it is necessary to exclude the structurally stable and non-deformable α-helix, β-sheet, glycosylation sites which may obscure the epitopes or alter the antigenicity, etc [8]. Even so, the predicted epitopes are still inaccurate [4]. Compared with the current study on SARS-CoV-2, this work adopted various prediction methods and 3D structure databases developed in recent years, which were based on artificial neural network, Hidden Markov Model(HMM), Support Vector Machine(SVM), etc, such as ABCpred, BepiPred2.0, SEPPA 3.0, IEDB, etc. Compared with prediction by a single method [9] or on the basis of epitopes of SARS [10], these methods and databases greatly improved the accuracy of prediction and had more bioinformatic meaning. We comprehensively analyzed the prediction results from the tools which were widely used, set up screening criteria on the basis of primary structure, secondary structure and tertiary structure, so that the prediction results would more accurate and reliable.

The S protein, the E protein and the M protein are surface proteins of SARS-CoV-2, which have the potential as antigenic molecules. However, the current study on the epitopes prediction of SARS-CoV-2 [11], due to the S protein has been reported to be the directly binding molecule of SARS-CoV-2 to ACE2[12], the prediction of epitopes is mainly focusing on the S protein, with few studies on the E protein and the M protein. In this work, we analyzed the S protein, the E protein and the M protein, predicted their epitopes. On the basis, 7 B cell epitopes were predicted, including 2 conformational and 6 linear B cell epitopes, one of the conformational and one of the linear are coincide. All of the epitope A, B, C, D located on the surface of the tail of the S protein, which is relatively easy to bind. The epitope E is located on the head of the S protein, which is the key area where the S protein recognizes and binds to ACE2 [12,13], has the potential to block the infection process. The epitope F and the epitope G located on the end of the head of the E protein, the two epitopes coincide, this may due to they are all consecutive and the secondary structure avoided the α-helix and the β-sheet. The epitope H is derived from the M protein, the structure and conservation could not be determined due to the inability to predict reliable structure. However, it could be known from the surface probablity scores that the epitope H is more likely to be located on the surface of the M protein.

The higher the conservation score calculated by the Consurf Server is, the more likely the site is to be mutated in the evolutionary process. When the score<1, the site is likely to be a conservative site; when the score is between 1 and 2, the site is a site which is likely to be a relatively easy mutation; when the score>2, the site is likely to be an easy mutation site [14]. In the 7 epitopes obtained, all the epitopes of the S, E, M protein were absolute conservative among all SARS-CoV-2 sequences. For the human coronavirus dataset and the coronavirus dataset, only the average conservative score of the epitope D is higher than 1, which is prone to mutation. The epitope D should not be used as an epitope of the S protein. The conservation of the epitope H could not be calculated by the PDB file, the application value of the epitope H needed further experimental verification. Although the epitopes could be integrally considered to be conservative, the independent residues of these epitopes could still easy to mutate. Except the epitope E, all of 6 dominate epitopes contain 1-2 residues which has a conservative score higher than 1(Table 3C), indicating that these residues were likely to be easy mutation sites. These residues mostly located at the head or the tail of the epitopes, therefore, the mutation of these residues should be paid attention to, and the length of the epitopes should be adjusted according to the actual effect in application. The scores of epitopes in different datasets were different, which could due to the quantity of sequences in the datasets and the structures were analyzed in different situations.

In this work, we predicted 6 reliable epitopes: A, B, C, E, F/G and H. The reliability of the epitopes of the S protein was relatively better than that of the epitopes of the E protein and the M protein, indicating that the S protein is still the optimal choice for the prediction of epitopes and the development of vaccine. All of the 6 epitopes were able to achieve absolute conservation in SARS-CoV-2, and to achieve relative conservation in the data set, including SARS, etc. Therefore, the epitopes not only have the potential to be directly applied on the treatment in this disease, but also have the potential to prevent the possible threats caused by other types of coronavirus. In addition, although various factors of prediction were integrated in this work, more experimental data are needed to further verify whether all the 6 epitopes can induce the body to produce corresponding antibodies and generate specific humoral immunity, due to the limited data set and other factors.

## Materials and methods

### Materials

All of the analysis was based on the NCBI Reference Sequence: NC_045512.2. We obtained the sequence of S, E and M protein and its proximal sequences by BLAST, which got 420, 334 and 329 sequences in total from NCBI database respectively. We obtained the whole genome sequence of SARS-CoV-2 from Genbank and GISAID(959 in total), which were used to be a dataset after genome annotation. The genome sequences, which performed mistakes of translation, were deleted.

### Methods

#### Basic analysis of surface protein of SARS-CoV-2

The physical and chemical properties of target protein were analyzed by the Port-Param tool in ExPASy [15], including the primary structure of the target protein, molecular formula, theoretical isoelectric point, the protein instability index(the index<40 means the protein was stable), *etc*. Online software, ProtScale, was used to deeply analyze the hydrophilicity and hydrophobicity of target protein and the distribution of hydrophilicity and hydrophobicity of polypeptide chains [15]. SARS-CoV-2 carried the S/E/M protein through the virus envelope, the transmembrane region of the protein was predicted online by TMHMM 2.0 [16].

#### Prediction of the 3D structure of target protein

With the amino acid sequences of the surface protein of SARS-CoV-2 of NC_045512.2 as templates, based on homology modeling method, we predicted the 3D structure through the online server SWISS-MODEL[17], selected and optimized the optimal structure based on the template identity and GMQE value[17], the rationality of the structure was evaluated by Ramachandran plot [18] with PDBsum server. The structures were displayed and analyzed by SWISS-pdb Viewer v4.10 [19].

#### Prediction of conformational B cell epitopes of target protein of SARS-CoV-2

Based on the structures, the conformational B cell epitopes were predicted by SEPPA 3.0 [20] and Ellipro [21] respectively, and the common predicted conformational B cell epitopes from two methods were selected for the further analysis.

#### Prediction of linear B cell epitopes of target protein of SARS-CoV-2

The Protean module of DNAStar was used to predict the flexibility[21], surface probablity [22] and antigenic index [23] of the target protein of SARS-CoV-2. The linear B cell epitope was predicted by ABCpred [24] and BepiPred 2.0 [25] respectively and the common predicted linear B cell epitopes from two methods were selected for the further analysis. Coupled with the secondary structure, the tertiary structure and the glycosylation sites [26] *etc*, the linear B cell epitopes were finally determined.

#### Analysis of epitope conservation

Based on the PDB model and the multiple alignment result, we used the Consurf Server to analyze the conservation of amino acid sites of the epitopes online[27]. The conservation of epitopes on the surface protein of SARS-CoV-2 was analyzed by multiple alignment with MAFFT and Logo was drawn with Weblogo [28,29].

## Authors’ contribution

JL conceived the study and participated in its design and coordination. HD participated in the design of the study and helped draft the manuscript. YB participated in analysis of conservation, sequence alignment and manuscript drafting. BZ participated in antigenic prediction. FC participated in drafting the manuscript. All authors read and approved the final manuscript.

## Acknowledgements

This work was supported by the National Key R&D Program of China (2018YFC0910201), the Key R&D Program of Guangdong Province (2019B020226001), and the Science and the Technology Planning Project of Guangzhou (201704020176). The Student Entrepreneurship and Innovation Center of the school of biology and biological engineering, South China University of Technology, also provided a lot of help during the preparation of the project.

## Supplementary information

**Figure S1** Deep analysis of hydrophilicity and hydrophobicity of surface protein of SARS-CoV-2

The online software, ProtScale, was used to predict the hydrophilicity and hydrophobicity of the surface protein deeply. **A.** The S protein has a maximum score of hydrophobicity, 3.222 at the 7^th^ site, which revealed a strong hydrophobicity; a minimum score of hydrophobicity, −2.589 at the 679^th^ site, which revealed a strong hydrophilicity. The score of hydrophilicity and hydrophobicity on the polypeptide chain of S protein constantly fluctuates, with most of the scores being negative, which revealed the possibility that the protein had bisexual properties on the basis of hydrophilicity. **B.** The E protein has a maximum score of hydrophobicity, 3.489 at the 21^st^ and the 25^th^ site, which revealed a strong hydrophobicity; a minimum score of hydrophobicity, −1.550 at the 65^th^ site, which revealed a strong hydrophilicity. Most of the scores of the residues being positive, which revealed the possibility that the protein has obvious hydrophobicity. **C.** The M protein has a maximum score of hydrophobicity, 2.978 at the 84^th^ site, which revealed a strong hydrophobicity; a minimum score of hydrophobicity, −1.956 at the 211^th^ and the 212^th^ site, which revealed a strong hydrophilicity. The scores of hydrophilicity and hydrophobicity on the polypeptide chain of M protein showed large fluctuations, and the number of positive scores and negative scores were similar, the positive scores accounted for the majority, which revealed the possibility that the protein had bisexual properties on the basis of hydrophobicity.

**Figure S2** The transmembrane region of the surface protein of SARS-CoV-2 The S, E and M protein are embedded in the envelope of SARS-CoV-2, the transmembrane helix was predicted by TMHMM 2.0 server. All of three amino acid indexes were higher than 18, indicating the reliability of the prediction. **A.** For the S protein, an outside-in transmembrane helix was predicted in the 23 residues of amino acids from position 1214^th^ to position 1236^th^ at the N-terminal. The amino acid index was 23.97303. **B.** For the E protein, an inside-out transmembrane helix was predicted in the 23 residues of amino acids from position 12^th^ to position 34^th^ at the N-terminal. The amino acid index was 25.72521. **C.** For the M protein, 2 outside-in transmembrane helices were predicted, which were a helix in the 20 residues of amino acids from position 20^th^ to position 39^th^ and a helix in the 23 residues of amino acids from position 78^th^ to position 100^th^ at the N-terminal. An inside-out helix was predicted in the 23 residues of amino acids from position 51^st^ to position 73^rd^ at the N-terminal. The amino acid index was 64.90522.

**Figure S3** The antigenic conservation of the surface protein in SARS-CoV-2 **A.** The epitope A was absolutely conservative in 756 SARS-CoV-2 genomes. **B.** The epitope B was absolutely conservative in 756 SARS-CoV-2 genomes. **C.** The epitope C was absolutely conservative in 756 SARS-CoV-2 genomes. **D.** The epitope D was absolutely conservative in 756 SARS-CoV-2 genomes. **E.** The epitope E was absolutely conservative in 756 SARS-CoV-2 genomes. **F.** The epitope F/G was absolutely conservative in 939 SARS-CoV-2 genomes. **G.** The epitope H was absolutely conservative in 913 SARS-CoV-2 genomes.

**Figure S4** The antigenic conservation of the surface protein in human coronavirus **A.** The conservation of the epitope A in 331 human coronavirus genomes. **B.** The conservation of the epitope B in 331 human coronavirus genomes. **C.** The conservation of the epitope C in 331 human coronavirus genomes. **D.** The conservation of the epitope D in 331 human coronavirus genomes. **E.** The conservation of the epitope E in 331 human coronavirus genomes. **F.** The conservation of the epitope F/G in 268 human coronavirus genomes. **G.** The conservation of the epitope H in 268 human coronavirus genomes.

**Figure S5** The antigenic conservation of the surface protein in coronavirus **A.** The conservation of the epitope A in 403 human coronavirus genomes. **B.** The conservation of the epitope B in 403 human coronavirus genomes. **C.** The conservation of the epitope C in 403 human coronavirus genomes. **D.** The conservation of the epitope D in 403 human coronavirus genomes. **E.** The conservation of the epitope E in 403 human coronavirus genomes. **F.** The conservation of the epitope F/G in 334 human coronavirus genomes. **G.** The conservation of the epitope in 327 human coronavirus genomes.

**Table S1 Bepipred2.0 linear epitope prediction results**

**Table S2 ABCpred linear epitope prediction results**

**Table S3 Prediction results of conformational B cell epitopes of surface protein by Ellipo**

**Table S4 Prediction results of conformational B cell epitopes of surface protein by SEPPA3.0**

**Table S5 Conservation analysis of epitopes in different datasets by Consurf**

