## Supplementary figures and images for "Prediction and Evolution of B Cell Epitopes of Surface Protein in SARS-CoV-2"

### Figure S1 Deep analysis of hydrophilicity and hydrophobicity of surface protein of SARS-CoV-2.pdf

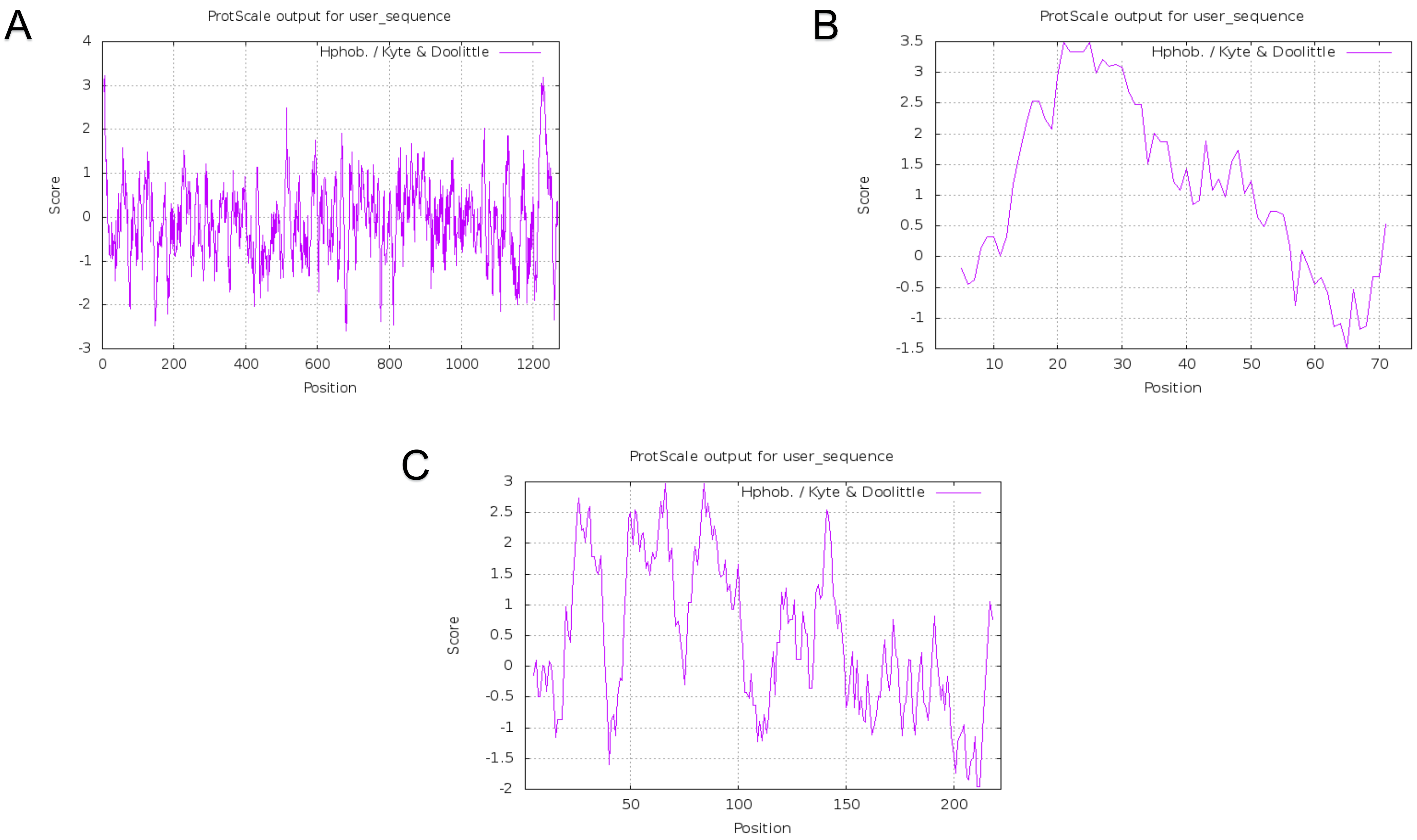

### Figure S2 The transmembrane region of the surface protein of SARS-CoV-2.pdf

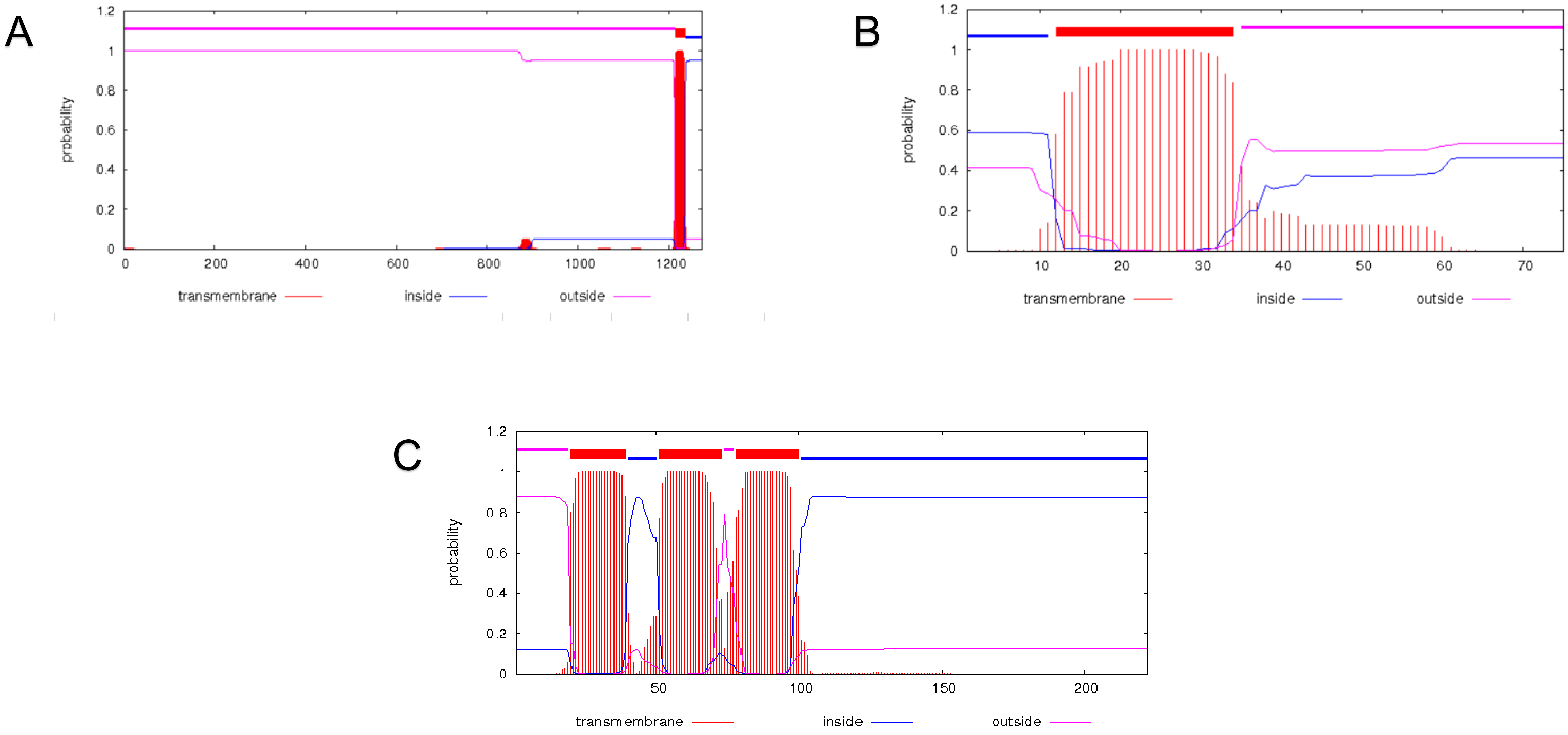

### Figure S3 The antigenic conservation of the surfaceS protein in SARS-CoV-2.pdf

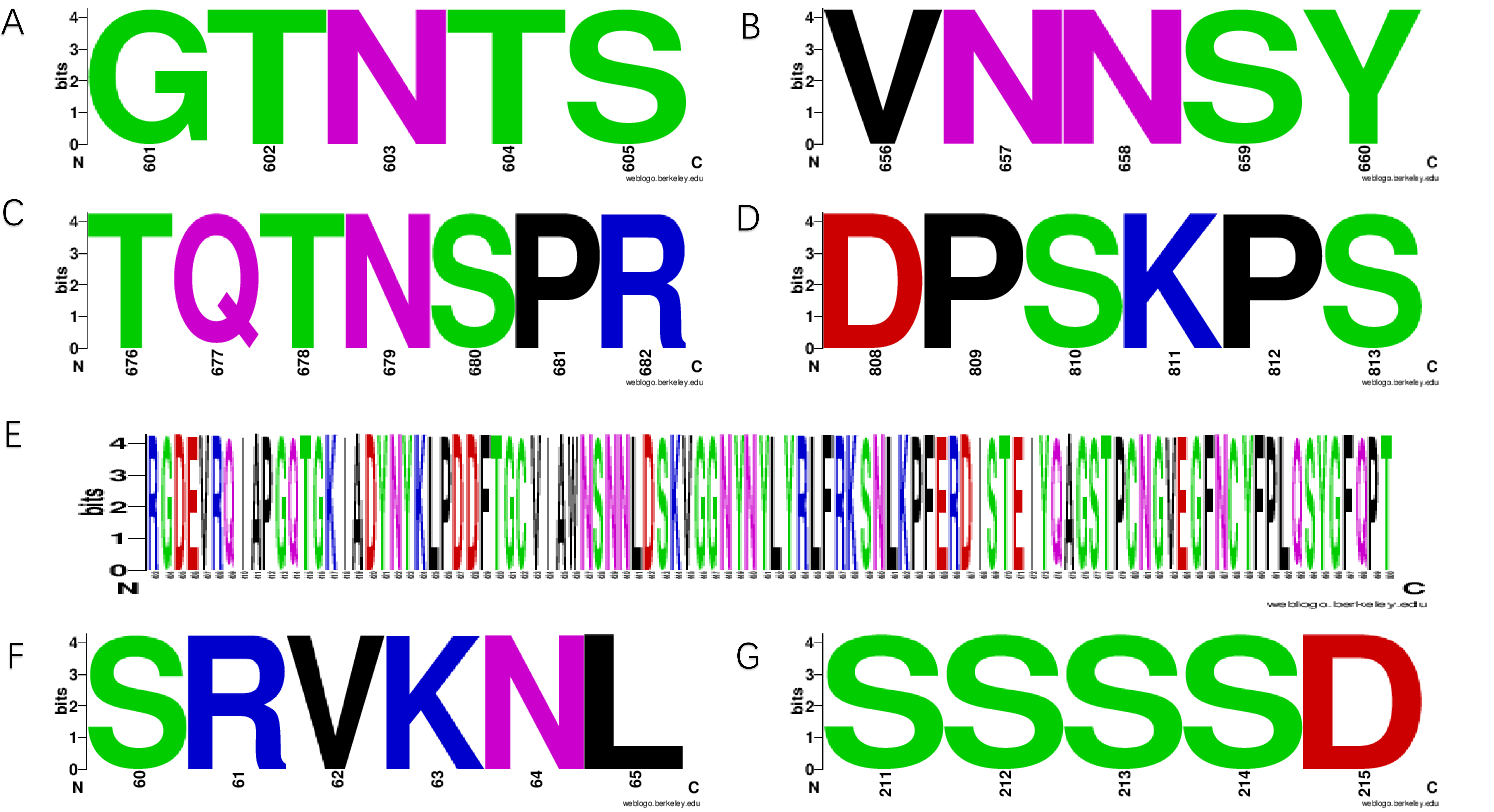

### Figure S4 The antigenic conservation of the surfaceS protein in human coronavirus.pdf

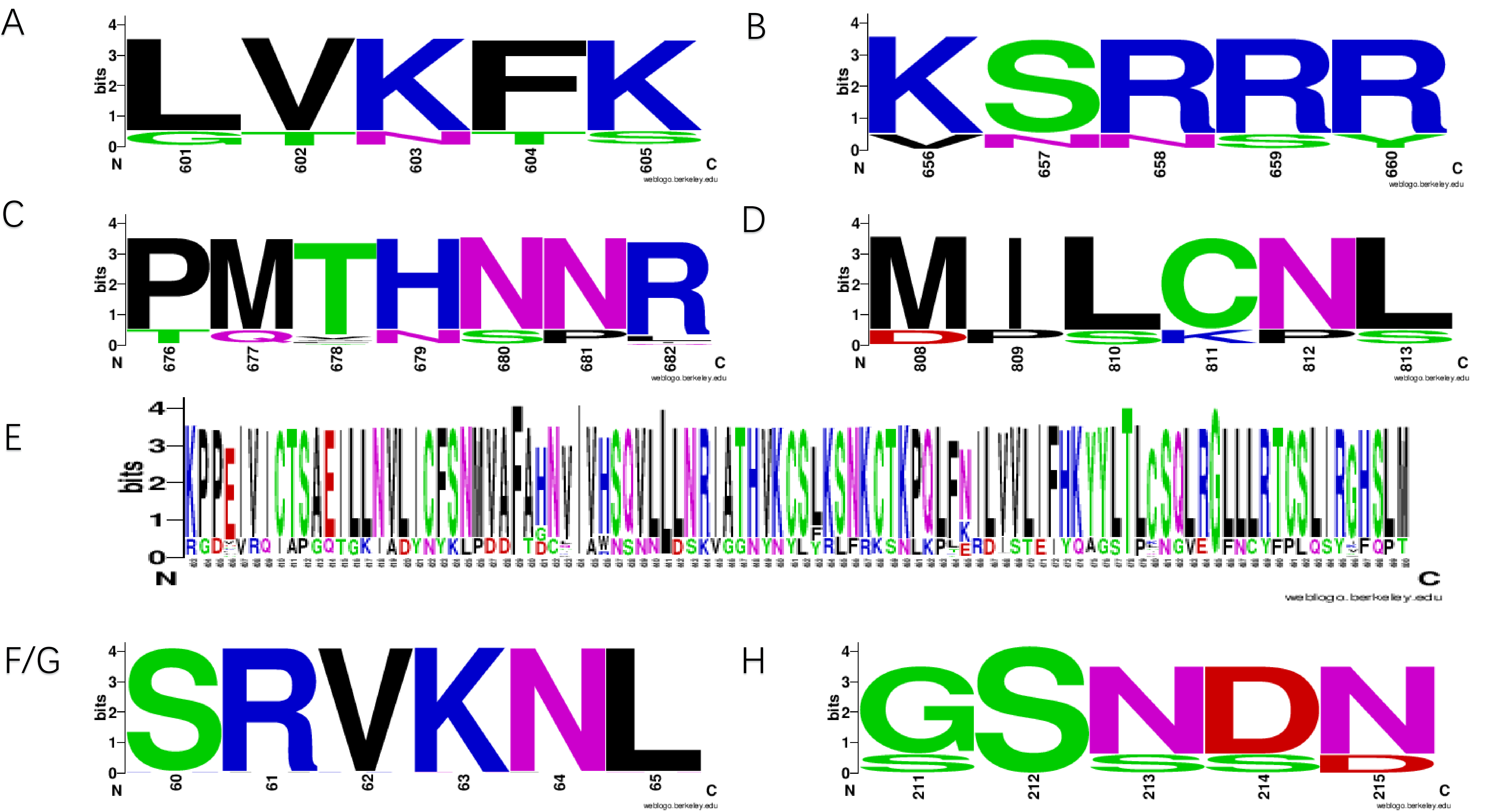

### Figure S5 The antigenic conservation of the surfaceS protein in coronavirus.pdf

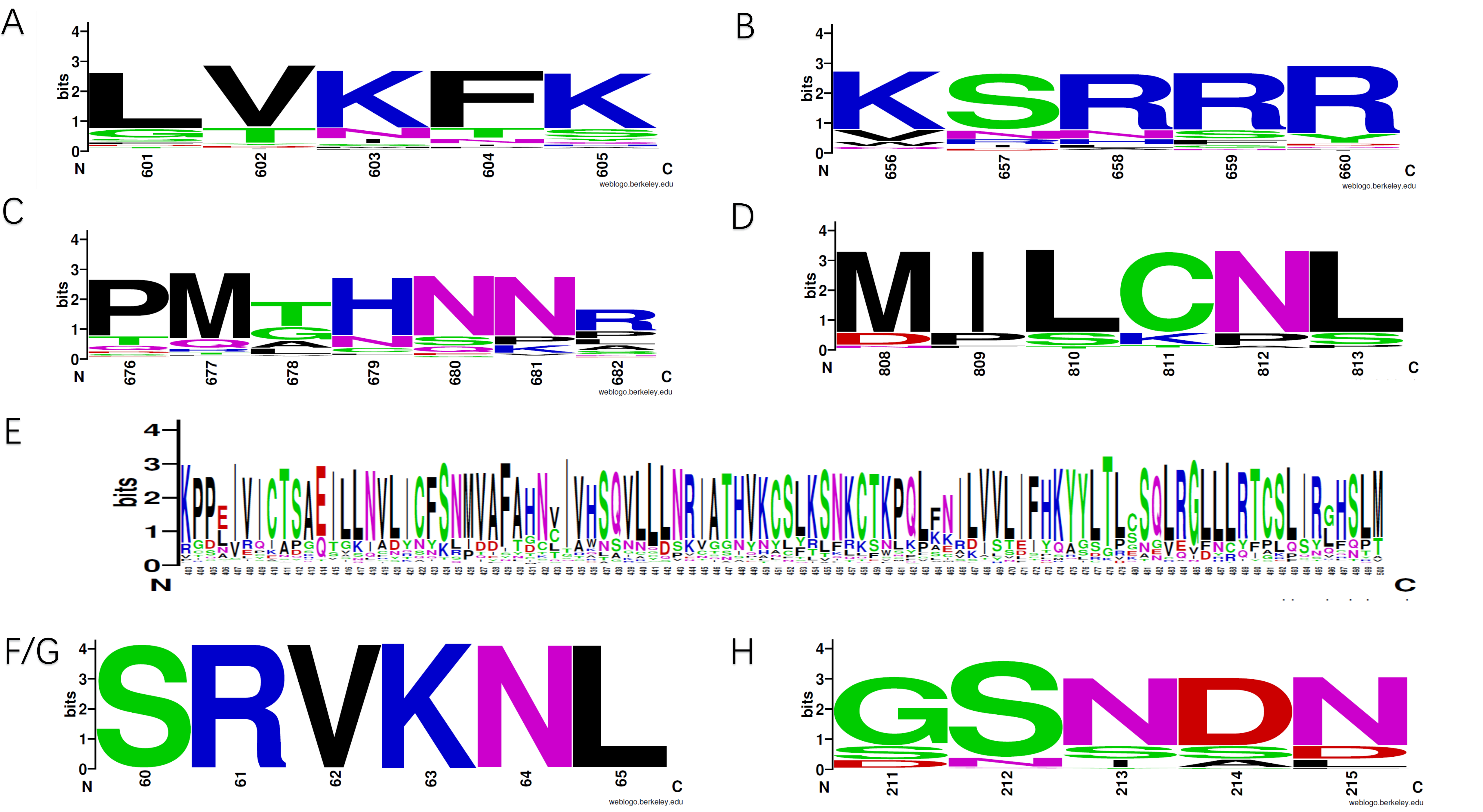
